# Spatial cell type mapping of multiple sclerosis lesions

**DOI:** 10.1101/2022.11.03.514906

**Authors:** Celia Lerma-Martin, Pau Badia-i-Mompel, Ricardo O. Ramirez Flores, Patricia Sekol, Annika Hofmann, Thomas Thäwel, Christian J. Riedl, Florian Wünnemann, Miguel A. Ibarra-Arellano, Tim Trobisch, Philipp Eisele, Denis Schapiro, Maximilian Haeussler, Simon Hametner, Julio Saez-Rodriguez, Lucas Schirmer

**Affiliations:** Division of Neuroimmunology, Department of Neurology, Medical Faculty Mannheim, Heidelberg University, Mannheim, Germany; Heidelberg University, Faculty of Medicine, and Heidelberg University Hospital, Institute for Computational Biomedicine, Bioquant, Heidelberg, Germany; Division of Neuropathology and Neurochemistry, Department of Neurology, Medical University of Vienna, Vienna, Austria; Institute of Pathology, Faculty of Medicine, Heidelberg University and Heidelberg University Hospital, Heidelberg, Germany; Genomics Institute, University of California, Santa Cruz, Santa Cruz, CA, USA; Mannheim Center for Translational Neuroscience, Medical Faculty Mannheim, Heidelberg University, Mannheim, Germany; Interdisciplinary Center for Neurosciences, Heidelberg University, Heidelberg, Germany

**Keywords:** single-nucleus RNA-sequencing, spatial transcriptomics, subcortical lesions, neuroinflammation

## Abstract

Multiple sclerosis (MS) is a prototypic chronic-inflammatory disease of the central nervous system. After initial lesion formation during active demyelination, inflammation is gradually compartmentalized and restricted to specific tissue areas such as the lesion rim in chronic-active lesions. However, the cell type-specific and spatially restricted drivers of chronic tissue damage and lesion expansion are not well understood. Here, we investigated the properties of subcortical white matter lesions by creating a cell type-specific spatial map of gene expression across various inflammatory lesion stages in MS. An integrated analysis of single-nucleus and spatial transcriptomics data enabled us to uncover patterns of glial, immune and stromal cell subtype diversity, as well as to identify cell-cell communication and signaling signatures across lesion and non-lesion tissue areas in MS. Our results provide insights into the conversion of the tissue microenvironment from a ‘homeostatic’ to a pathogenic or ‘dysfunctional’ state underlying lesion progression in MS. We expect that this study will help identify spatially resolved cell type-specific biomarkers and therapeutic targets for future interventional trials in MS.

## Introduction

Multiple sclerosis (MS) is a prototypic and the most common inflammatory disease of the central nervous system, characterized by multifocal demyelination, axonal damage and gliosis, ultimately leading to a loss of neurons across various gray matter areas^1–3^. A major problem associated with progressive disease in MS is compartmentalized inflammation^4^.

An important anatomical niche of compartmentalized inflammation in MS is formed by inflammatory aggregates in deep meningeal sulci resembling ectopic lymphoid follicle-like structures. It is widely accepted that sustained meningeal inflammation is a key factor contributing to cortical pathology and, particularly, resulting in neuronal injury of upper cortical layers^5–7^. Another important niche area of chronic tissue damage and sustained inflammation in MS are subcortical white matter lesion rims (LRs)^8^. Several postmortem and in vivo magnetic resonance imaging studies have demonstrated that these chronic active lesions are regularly accompanied by iron uptake at LRs, centrifugal lesion expansion and progressive disease^9–12^. A comprehensive understanding of the cellular composition and molecular dynamics of the lesion tissue microenvironment would have a critical impact on future biomarker and interventional studies in individuals diagnosed with MS.

MS lesions follow a well-characterized temporal sequence of the level and spatial pattern of inflammation and tissue damage, thus providing information about the lesion age^13–15^. Acute MS lesions (MS-A) commence through active myelin breakdown with presence of myelin-phagocytosing cells including macrophages as well as activated microglia and astroglial subtypes^16–18^. After this initial stage, lesions may either remyelinate^19^ or enter a chronic active stage (MS-CA) with a distinct inflammatory LR and a well-demarcated completely demyelinated lesion core^9,10^. Eventually, MS-CA lesions convert into an inactive stage (MS-CI) without LR inflammation but low-level microglial activation and a dense astroglial scar meshwork throughout the lesion^20,21^. Nevertheless, the precise lesion kinetics are not well understood, and it is known that chronically inflamed and iron-positive LRs can persist for years in certain individuals with MS^8,22^. Also, chronic MS lesions may convert into remyelinated lesions representing myelin repair in at least a subset of individuals^23^. Remyelination might also remain incomplete and is then often seen adjacent to LR areas^24^. Over the past decades, histopathological and MR imaging studies have collectively provided deep insights into inflammatory lesion development and temporal lesion dynamics in individuals diagnosed with MS^25,26^. However, due to a lack of spatially resolved molecular tools it has not been possible to discern spatially restricted areas of cell type-specific pathology and eventually map back these changes to defined lesion and non-lesion tissue niches.

Here, we utilized single-nucleus RNA-sequencing (snRNA-seq) paired with spatial RNA-seq (spatial transcriptomics, ST) to create a large-area map of gene expression covering lesion core and LR areas as well as normal-appearing white matter. Using a wide range of computational tools, we estimated the distribution of cell types across subcortical MS lesion areas, characterized tissue niches and identified spatially restricted cell-cell interaction and signaling pathway signatures. To facilitate the use of these results by the community, we have built an interactive single-cell and ST web browser to visualize MS lesion transcriptomics. Our spatially resolved gene expression underlying MS lesion development should help facilitate the development of region- and cell type-specific biomarkers and design targeted treatments to tackle compartmentalized inflammation in MS.

## Results

### Transcriptomic profiling of human multiple sclerosis in WM

To unravel the transcriptomic signatures linked to the spatial environment in MS pathology, we characterized 7 control (CTRL) and 21 MS tissue samples. In detail, we focused on subcortical white matter tissues and performed ST on 17 tissue blocks (4 CTRL vs 13 MS) with paired snRNA-seq on the same blocks (3 CTRL vs 8 MS) (Methods) (**Fig. 1a and Supplementary Tables 1-2**). Based on the inflammatory lesion activity, we classified MS lesions into acute active-demyelinating (MS-A), chronic active (MS-CA) and chronic inactive (MS-CI) stages (Methods) (**Fig. 1b)**. For the paired snRNA-seq dataset, we obtained a total of 69,526 nuclei (n=11, average=6,321) with a mean of 2,042 detected genes per nucleus after filtering out low-quality profiles and potential doublets (**Extended Data Fig. 1a**). After controlling for data quality, the ST dataset contained a total of 52,945 spots (n=17, average = 3,114) and a mean of 1,224 genes per spot (**Extended Data Fig. 1b, d**).

**Fig. 1.**
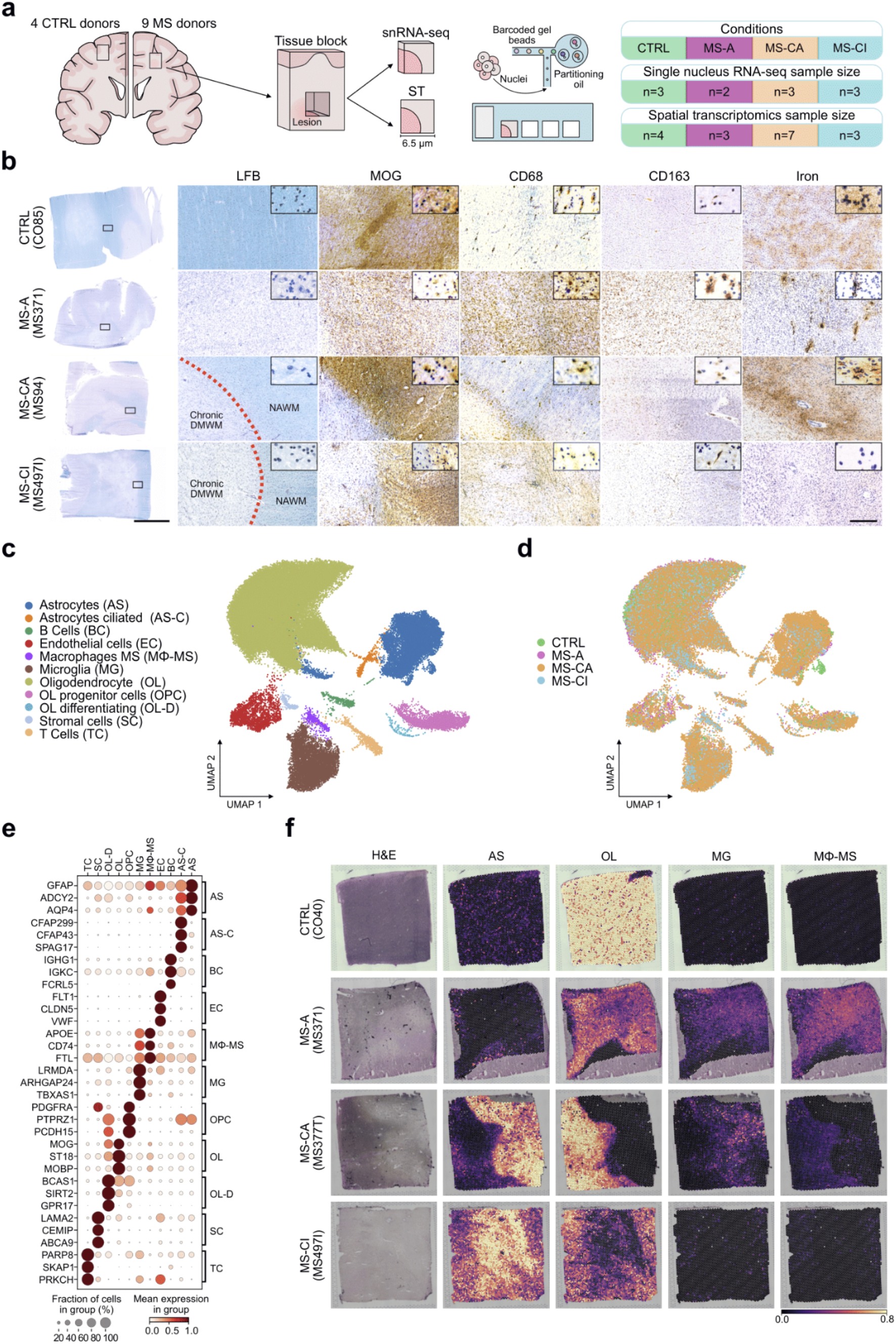
Spatial and cell type profiling of subcortical control and MS tissues. **a**, Study design showing different data modalities. CTRL, control tissue. MS-A, acute MS lesion type tissue. MS-CA, chronic-active MS lesion type tissue. MS-CI, chronic-inactive MS lesion type tissue. **b**, Control and MS tissue assessment by LFB and iron histology as well as immunohistochemistry for MOG (myelin), CD68 (activated myeloid cells) and CD163 (iron-metabolizing myeloid cells) marker proteins. Red dashed line indicates legion rim (LR) area. DMWM, demyelinated white matter. NAWM, normal appearing white matter. Overview images: scale bar 1 cm; zoom-in images: scale bar 500 µm. **c**, UMAP of snRNA-seq data from all samples (n = 69,526), color corresponds to annotated cell types. **d**, UMAP of snRNA-seq data, color corresponds to MS lesion type of samples. **e**, Dotplot of averaged z-transformed gene expression for marker genes. **f**, Characterization of ST data of control and MS lesion types using cell type deconvolution.

Based on snRNA-seq, we established an atlas comprising all major cell types present in human subcortical white matter tissues of both control and MS donors (**Fig. 1c, d**). After batch correction (Methods), we could identify five major subcortical cell types including astrocytes (AS), endothelial cells (EC), microglia (MG), oligodendrocyte progenitor cells (OPC), oligodendrocytes (OL) as well as six disease-enriched clusters including a new AS subtype expressing marker genes associated with primary cilia formation (*CFAP299, CFAP43* and *SPAG17*) that we annotated as ciliated astrocytes (AS-C). Other MS-enriched cell types were annotated as differentiating oligodendrocytes (OL-D), B cells (BC), macrophages (MΦ-MS), stromal cells (SC) and T cells (TC) (**Fig. 1e**). To corroborate our cell type annotations, we compared their transcriptional profiles to the ones from another recent study^8^ on human subcortical WM MS lesions and observed a high overlap between them (Methods) (**Extended Data Fig. 1c**).

Since ST captures multiple cells per spot, we increased its resolution by inferring cell type composition per spot (Methods). Using the paired snRNA-seq atlas as reference, we deconvoluted the ST tissue slides and found that mapped cell types matched the expected tissue architecture across controls and MS lesion types (**Fig 1f**).

A high abundance of AS was mapped to chronic active and inactive demyelinated white matter (DMWM), representing the astroglial scar tissue^27^. Likewise, spatial distribution of OL cells across different MS lesion types was mainly restricted to normal-appearing white matter (NAWM), as expected. Based on transcriptomic signatures, we could map MG and MΦ-MS cells to inflamed MS-A lesion core and MS-CA LR areas. Also, we could only map a few MG and barely any MΦ-MS profiles to control and MS-CI tissues. In MS-CA lesion areas, we additionally found a strong compartmentalization of both MG and AS cells restricted to DMWM areas. As compared to MG, MΦ-MS cells mapped to the rim of chronic active lesion areas with a characteristic high iron uptake^28^ (**Fig. 1b, e**).

The resulting inferred cell type compositions matched well the cell type proportions observed by snRNA-seq, both at the sample and cell type level (**Extended Data Fig. 1e, f**). Collectively, integration of snRNA-seq with ST enables us to generate a comprehensive paired dataset of subcortical WM tissues enabling us to investigate expression profiles and regional relationships of glial, stromal, and immune cell types. To explore both snRNA-seq and ST data sets in an interactive and user-friendly environment, we have provided all data in a web browser (Data and code availability).

### Spatial organization of subcortical tissue

Next, we explored the spatial organization of the subcortical tissue microenvironment by leveraging the ST data. By unsupervised clustering of spots from all tissue sections, we identified seven clusters that could be linked to major cell types and specific ST tissue niches (Methods) (**Fig. 2a and Extended Data Fig. 2a**). Of note, the obtained ST niche clustering provides a comprehensive look into the structural transcriptomic assembly of the tissue blocks across controls and MS samples (**Extended Data Fig. 2b**).

**Fig. 2.**
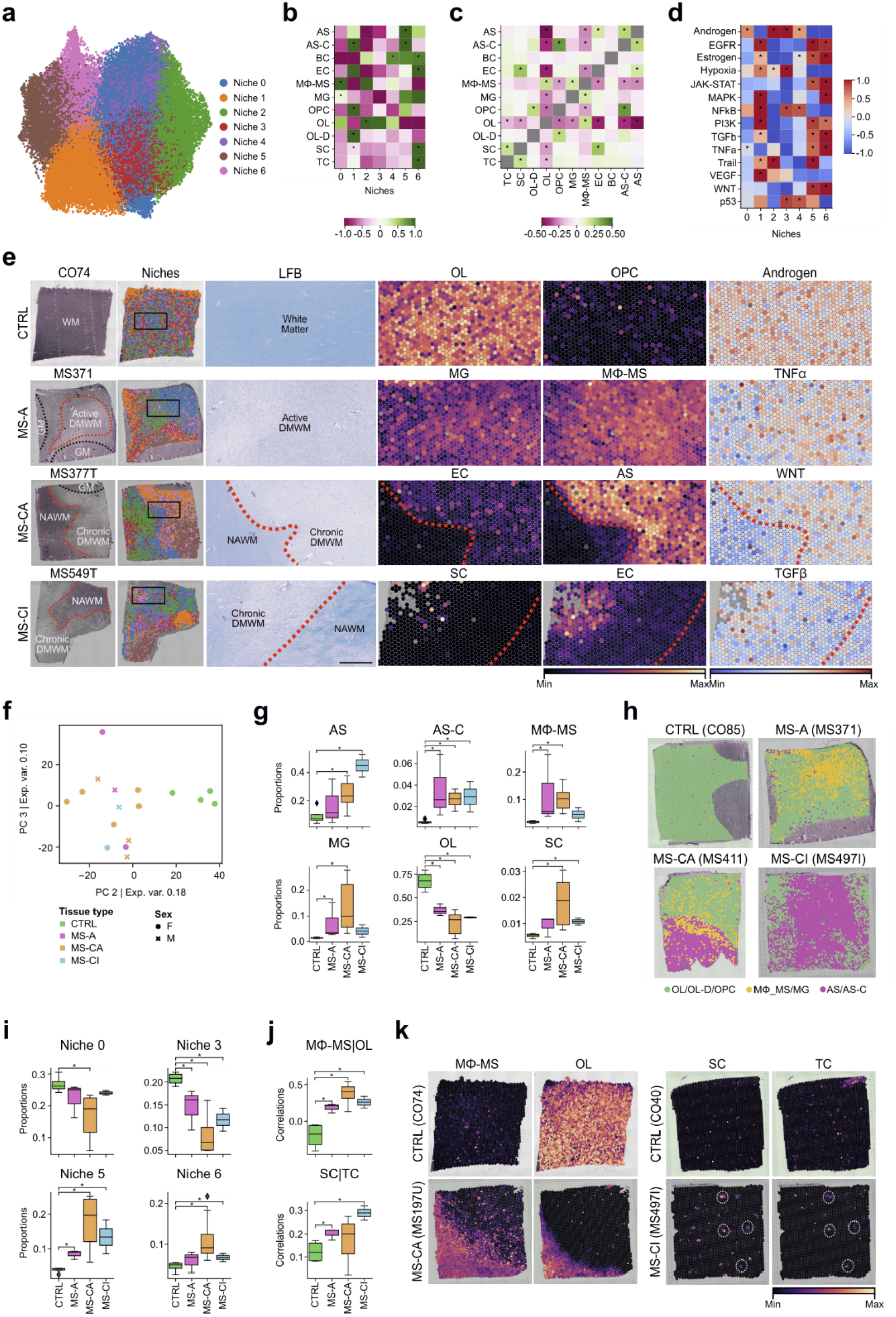
Spatial organization of subcortical control and MS tissues. **a**, Cell type niches UMAP of ST spots based on cell-type composition. Color indicates assigned niches labels by clustering. **b**, Scaled mean cell type composition per tissue niche. Asterisks indicate increased composition of a cell type per niche compared with other niches (one-sided Wilcoxon rank-sum test, adjusted P < 0.05). **c**, Mean Pearson correlation between cell type composition across spatial spots including all tissue samples. Asterisks indicate significant mean correlations (abs(mean corr) > 0.1 and mean P < 0.1). **d**, Mean scaled pathway activities within each niche. Asterisks indicate increased activity of a pathway in a niche compared with other niches (one-sided Wilcoxon rank-sum test, adjusted P < 0.05). **e**, Representative samples of control and MS lesion type tissues (panels left to right): HE histology, compositional tissue niches, LFB histology: scale bar 1.25 mm, cell type tissue mapping (two panels), pathway activity tissue mapping. Red dashed line indicates LR and black dashed line indicates cortical gray matter areas. WM, white matter. GM, gray matter. **f**, PCA projection of ST profiles. Color indicates lesion type and shape indicates gender. **g**, Boxplots of deconvoluted cell type in ST tissue slides. Color indicates lesion type, and asterisks significant differences between groups. **h**, Examples of tissue mapping for major glial cell types in ST tissue slides. Green indicates oligodendrocyte lineage cells (OL, OL-D and OPC), yellow indicates myeloid cells (MΦ-MS and MG) and pink astrocyte subtypes (AS and AS-C). **i**, Boxplots of niche compositions across lesion types. Color indicates lesion types, and asterisks significant differences between groups. **j**, Boxplots of co-localization correlations between cell types. Color indicates lesion types and asterisks significance between groups **k**, Examples of cell-type colocalization events in ST slides. Color indicates cell type abundance. Wilcoxon rank-sum test, adjusted P < 0.1 (**g, i, j**).

To further characterize ST niches, we investigated cell type enrichment within each tissue niche when compared to other niches (Methods) (**Fig. 2b**). Here, we found two main groups of cell type niches. The first group (0, 2, 3 and 4) appeared to be enriched by OL cells and was restricted to NAWM and MS LR areas (**Extended Data Fig. 2b**). The second group (1, 5 and 6) was enriched by AS and associated with DMWM areas. Niche 0 had increased numbers of OL, MΦ-MS and MG cell types and,mapped to inflamed lesion core and LR areas in MS-A and MS-CA typed tissues (see also **Fig. 2i**). Niche 1 was enriched by AS and cell types like OPC and OL-D. Niche 5 was overrepresented by AS subtypes including AS-C cells. Finally, tissue niche 6 was characterized by cell type enrichment of AS, TC, BC, SC and EC subtypes and histologically mapped to areas with increased blood vessel density, suggesting a perivascular tissue niche associated with immune cell infiltration (**Fig. 2b, Fig. 2i and Extended Data Fig. 1d, 2b**). These results provide a wide description of cell type organization across all tissue slides.

To complement the identification of specific colocalization events based on tissue niche analysis, we next tested correlations between compositions of cell types in the same spatial spots across samples and lesion stages (Methods) (**Fig. 2c**). We found colocalizations between AS-C, OPC and AS, and between SC, EC, TC and AS cells. These colocalization events overlapped with the enrichment of these cell types in niches 1 and 6, respectively. A colocalization between OL, MG and MΦ-MS in niche 0 was also identified, suggesting potential engagement of myeloid cells in myelin breakdown.

To link these structural building blocks to molecular functions in tissues, we further described them by inferring signaling pathway activities at spot level across slides and lesions (Methods) (**Fig. 2d**). Niches enriched by OL (0, 2, 3 and 4) showed increased activity of androgen-related pathways (one-sided Wilcoxon rank-sum test, adjusted P < 0.05), independent of gender (**Extended Data Fig. 2c**). Tissue niches overrepresented by AS (1, 5 and 6) presented higher activity for pathways involved in tissue remodeling and maintenance including PI3K and TGFβ signaling (one-sided Wilcoxon rank-sum test, adjusted P < 0.05). Additionally, niches 5 and 6 showed enrichment for proinflammatory pathways such as TNFα and JAK-STAT (one-sided Wilcoxon rank-sum test, adjusted P < 0.05), suggesting roles in compartmentalized tissue inflammation.

We found that control samples did not show spatial tissue segregation, indicating a homogenous tissue niche composition with different glial cell types ‘stochastically’ distributed throughout the tissue section (**Extended Data Fig. 2a-b**). OL cells were the main cell type in control tissues and showed spatial enrichment for genes associated with androgen signaling such as *NDRG1* and *DBI* (**Fig. 2e**). In lesion areas of MS-A samples, we found that MΦ-MS and MG cells were associated with TNFα signaling, a classic proinflammatory pathway in chronic inflammation (**Fig. 2e**). In MS-CA lesion areas, the inflamed LR formed a distinct spatial niche that separated NAWM from DMWM lesion core areas (**Fig. 2e**). In MS-CA typed tissues we noted that certain tissue niches appeared to be enriched for mesenchymal SC and EC cell types linked to WNT and TGFβ signaling, as seen in a previous study^29^.

Overall, these results suggest that the MS tissue microenvironment is characterized by a dynamic cell type and tissue niche patterning driven by glial subtype-associated signaling pathways playing critical roles in tissue inflammation and remodeling.

### Structural variation of subcortical tissue

Once we analyzed the tissue architecture across slides and conditions, we explored the differences between control and MS lesion type tissues. Principal component analysis (PCA) was computed to unbiasedly identify the main sources of variability based on pseudo-bulk gene expression profiles of ST tissue sections (Methods) (**Fig. 2f and Extended Data Fig. 2d**). Of note, control samples clustered separately from MS ones. While we could not distinguish different MS lesion types, we observed a change in cell proportion and their spatial distribution in both ST and snRNA-seq datasets (**Fig. 2g, h, Extended Data 2d, e and Supplementary Table 3**). As predicted, numbers of OL cells decreased while AS subtypes increased during lesion development (Wilcoxon rank-sum test, adjusted P < 0.1). In acute tissue samples, we noted a high density of inflammatory cells such as MΦ-MS, TC and BC subtypes; as well as activated MG with LR mapping of these cells in MS-CA tissues (Wilcoxon rank-sum test, adjusted P < 0.1). Further, as predicted from histological assessment we found that AS cells localized to MS-CA and MS-CI DMWM areas. AS-C were overrepresented in all lesion types compared to control tissues (Wilcoxon rank-sum test, adjusted P < 0.1), providing additional insight into this ciliated and, potentially, functionally distinct scar tissue AS subtype (**Fig. 2g and Extended Data 2f**).

To further characterize changes in proportions across different conditions, we examined tissue niches as described above (**Fig. 2i, Extended Data 2f, g and Supplementary Table 4**).

Indeed, niche 3, resembling a ‘homeostatic’ NAWM signature, was more enriched in control than in MS tissues (Wilcoxon rank-sum test, adjusted P < 0.1). Of note, niche 0 changed when comparing control to MS tissues and could resemble a tissue niche associated with active demyelination in early MS-A tissue due to a compositional abundance of MΦ-MS and activated MG (Wilcoxon rank-sum test, adjusted P < 0.1) (see also **Fig. 2b**). Tissue niches 5 and 6 were enriched in MS-CA relative to control tissues (Wilcoxon rank-sum test, adjusted P < 0.1), in agreement to their proinflammatory tissue profile. Here, we found a colocalization of TC and SC cell types, hence likely resembling perivascular immune cell cuffing in inflamed MS tissue areas as described above (**Fig. 2b, j, k, Extended Data 2h and Supplementary Table 5**).

By performing a thorough transcriptomic tissue analysis based on paired snRNA-seq and ST, we could identify tissue niche clusters that were linked to spatially restricted MS lesion tissue areas.

### Characterization of MG subpopulations

Next, we focused on the molecular heterogeneity of MG cells, which is central for lesion formation and tissue injury in both early MS-A^30^ and MS-CA lesions^8^. To this end, we subclustered MG nuclei based on their transcriptomic profile and found five distinct MG cell states (Methods) (**Fig. 3a and Extended Data 3a, b**). We then tested if the relative abundance of MG cells changed across lesion types (Wilcoxon rank-sum test, adjusted P < 0.1) (Methods). We observed that cell state 0 was overrepresented in control samples, suggesting that this MG subtype would be related to a homeostatic subtype strongly associated with NAWM tissue niches. Conversely, MG cell states 1, 2 and 3 were relatively more abundant in MS than in control samples, suggesting that these MG subtypes were linked to activated MG cells localized to inflamed MS lesion areas such as the LR (**Fig. 3b, Extended Data 3c and Supplementary Table 6**).

**Fig. 3.**
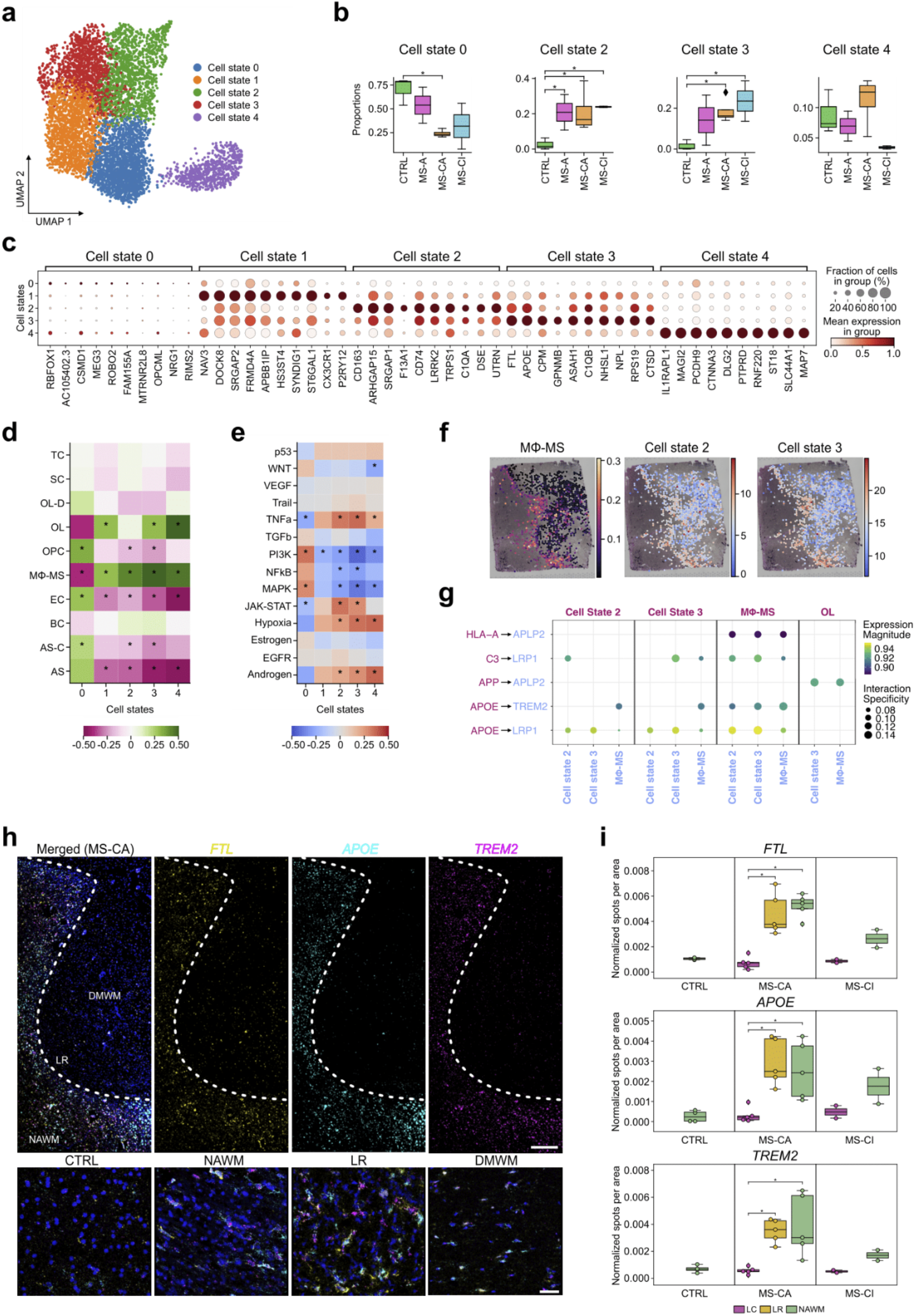
Identification and spatial characterization of MG cell states. **a**, UMAP of MG cell states based on snRNA-seq. Color indicates assigned MG subtype state. **b**, Boxplots of cell state compositions based on snRNA-seq. Color indicates MS lesion type and asterisks differences between groups (Wilcoxon rank-sum test, adjusted P < 0.15). **c**, Dotplot showing scaled mean expression of marker genes per MG cell state. **d**, Mean correlations between MG cell state scores and cell type composition across spots from MS-CA tissue slides. **e**, Mean correlation between MG cell state scores and pathway activities across ST spots from MS-CA tissue slides. Asterisks (d, e) indicate mean correlations (abs(mean corr) > 0.15 and mean P < 0.05). **f**, Example of colocalization of MΦ-MS composition with MG states 2 and 3 scores in a chronic active slide. Spots with less than 11.11% of MG were removed. **g**, Specific cell-cell ligand-receptor interactions inferred from snRNA-seq, arrow indicates direction of ligand-receptor signaling. **h**, smFISH of MS-CA tissue for *FTL, APOE* and *TREM2*. Overview images: scale bar 500 µm; zoom-in images: scale bar 40 µm. **i**, Boxplots of smFISH quantification across different tissue areas. Color indicates different lesion and non-lesion areas, asterisks indicate differences between groups (Wilcoxon rank-sum test, adjusted P < 0.05).

To better understand the obtained cell states, we next sought to identify marker genes per state (Methods). No specific marker genes could be obtained for MG cell state 0 relative to other MG clusters; however, we observed a certain enrichment of neuronal marker genes such as *RBFOX1* and *ROBO2* (**Fig. 3c and Supplementary Table 7**). Classical genes associated with homeostatic MG function were identified in cell state 1 (*DOCK8, P2RY12* and *CX3CR1*). MG cell state 2 showed enrichment for myeloid cell activation markers such as *CD163*, a scavenger receptor for hemoglobin-haptoglobin and iron uptake, *F13A1* encoding the coagulation factor XIII A subunit, and *CD74* encoding the HLA class II histocompatibility antigen gamma chain^31^. Cell state 3 was also enriched for genes linked to myeloid cell activation states and neuroinflammation such as *GPNMB* and, specifically, genes associated with lysosomal function and lipid metabolism such as *CTSD*, encoding cathepsin D, and *APOE*. In addition, MG cell states 2 and 3 were also characterized by upregulation of complement factor genes like *C1QA* (cell state 2) and *C1QB* (cell state 3)^8^. Further, we found that cell state 4 showed enrichment of OL marker genes (*ST18, RNF220* and *CTNNA3*) based on our current and previous snRNA-seq data^5^. Of note, marker genes in MG cell states 1 and 4 suggests a potential phagocytic role in clearance of neuronal (cell state 1) and OL cell lineage (cell state 4), material in line with our previous report on the identification of myelin-phagocytosing cells based on snRNA-seq^5^.

When comparing the transcriptional profiles of our myeloid cell types, including both MΦ-MS and MG, to a previous snRNA-seq dataset by Absinta *et al*^*8*^ (Methods), we confirmed overlap between our MG cell state 3 and the previously reported ‘foamy MΦ’ cluster. Further, our MΦ-MS cell type showed strong overlap with the described ‘iron MΦ’ cluster, and our MG cell state 4 showed overlap with the reported, however, not-annotated ‘state 4’ according to Absinta *et al* (Wilcoxon rank-sum test, adjusted P < 0.05) (**Extended Data Fig. 3d**).

As myeloid subtype cells play key roles in lesion formation and, specifically, LR inflammation, we next focused on chronic active MS lesions. After performing differential gene expression analysis comparing MG cells from MS-CA with control samples based on snRNA-seq (Methods) (**Extended Data Fig. 3e and Supplementary Table 8**), we observed an upregulation of proinflammatory marker genes associated with cell-cell communication (*CD84, CD14*) and gene regulation (*MAFB*)^32^. In addition, we noted downregulation of genes associated with anti-inflammatory and apoptosis-related cellular responses such as *INPP5D*^*33*^. The same trend could be retrieved from inferred pathway activities (Methods) that confirmed upregulation of proinflammatory signaling pathways like TNFα, NFκB and JAK-STAT and, conversely, downregulation of ‘homeostatic’ tissue maintenance pathways like TGFβ (**Extended Data Fig. 3f**). Transcription factor (TF) activity inference (Methods) corroborated this result since EGR1, one of the top regulators of TGFβ, was downregulated. Conversely, we noted that SPI1, also known as PU.1, an important developmental TF in MG cells^34^, and previously reported to be a genetic risk gene associated with MS^35^, was the most upregulated TF in MG cells derived from MS-CA tissues (**Extended Data Fig. 3g**).

Next, we mapped inferred MG states to tissue niches samples harboring MS-CA lesions (Methods). We found that the proinflammatory MG cell states 2 and 3 were positively correlated with OL and MΦ-MS cell types, but negatively with OPC, AS, AS-C and EC (P < 0.05) (Methods) (**Fig. 3d)**, suggesting roles in myeloid and OL cell interaction as expected at inflamed MS LRs with active demyelination. We also observed that MG cell states 2 and 3 showed overlap in signaling pathway activity (Methods) with upregulation of proinflammatory pathways (TNFα, JAK-STAT) and hypoxia-related pathways (**Fig. 3e)**. Additionally, enrichment of immune related gene sets was observed in MG cell states 2 and 3 (Methods) (**Extended Data Fig. 3h**). Further, cross-lesion stage comparison revealed that proinflammatory MG states 2 and 3 together with MΦ-MS cells mapped to MS-CA LR areas (**Fig. 3f**) and the lesion core in tissue samples with acute active-demyelinating lesions. This pattern could not be observed in control and chronic inactive lesion tissues (**Extended Data Fig. 3i**).

In summary, MG subclustering, cell state-specific gene expression and signaling pathway analysis helped identify functionally relevant subtypes and distinguish homeostatic from proinflammatory and potentially phagocytosing myeloid cell subtypes present at distinct lesion areas as shown by spatial mapping.

### Characterization of MG and MΦ-MS ligand-receptor interactions

Next, we investigated interactions of proinflammatory myeloid cells such as MΦ-MS and MG subtypes 2 and 3 with neighboring OL cells as the natural target cell type in MS (**Fig. 3g and Supplementary Table 9**). We inferred ligand-receptor interactions and focused on pairs of genes that were most correlated in MS-CA ST tissue slides (Methods). We identified the APOE-TREM2 relationship as an enriched ligand-receptor interaction pair between myeloid cells in MS-CA types tissues^8^, suggesting enhanced MG regulatory function including myelin debris clearing at LRs according to previous reports^36,37^. Similarly, we noted enhanced APOE-LRP1 signaling between MΦ-MS and MG subtypes, highlighting lipid and apoptotic cell clearance as key myeloid cell functions at inflamed MS LRs^38^. Further, APP-APLP2 ligand-receptor interaction was increased between OL cells (APP) and proinflammatory myeloid cells (APLP2) focusing on the LR, which also points towards regulatory functions of MΦ-MS and MG subtypes in MS^39,40^.

To validate enrichment of *APOE* and *TREM2* transcripts at MS LRs, we performed single molecule fluorescence *in situ* hybridization (smFISH) on MS-CA tissues. As MS-CA LRs show enhanced iron uptake, we also combined *FTL*, encoding ferritin light chain, to confirm mapping to iron-rich LRs (**Fig. 3h and Supplementary Table 10**). Of note, we observed increased smFISH signals for all three transcripts at both LR and NAWM areas with a trend towards even stronger expression of *TREM2* in NAWM (Methods). No increase of these marker genes was observed in control tissues and chronic inactive lesion areas (Wilcoxon rank-sum test, adjusted P < 0.05) (Methods) (**Fig. 3i and Supplementary Table 11**).

Collectively, ligand-receptor analysis demonstrated enrichment of key ligand-receptor interactions such as APOE-TREM2 with their transcripts mapping to both LR and NAWM areas.

### Characterization of AS subpopulations

In addition to myeloid cells, AS subtypes play critical roles in MS pathogenesis due to their ability to either help propagate inflammation or promote tissue homeostasis^31,41^. First, we decoded the transcriptomic heterogeneity of AS lineage cells and identified five different AS cell states (Methods) (**Fig. 4a and Extended Data Fig. 3j, k**). We then tested whether their relative abundances changed across MS lesion types (Methods) and found that AS cell states 2 and 3 had higher relative abundances in MS-CA relative to control tissues; further, AS cell state 2 was enriched in MS-CI tissues (Wilcoxon rank-sum test, adjusted P < 0.1) (**Fig. 4b and Supplementary Table 6**).

**Fig. 4.**
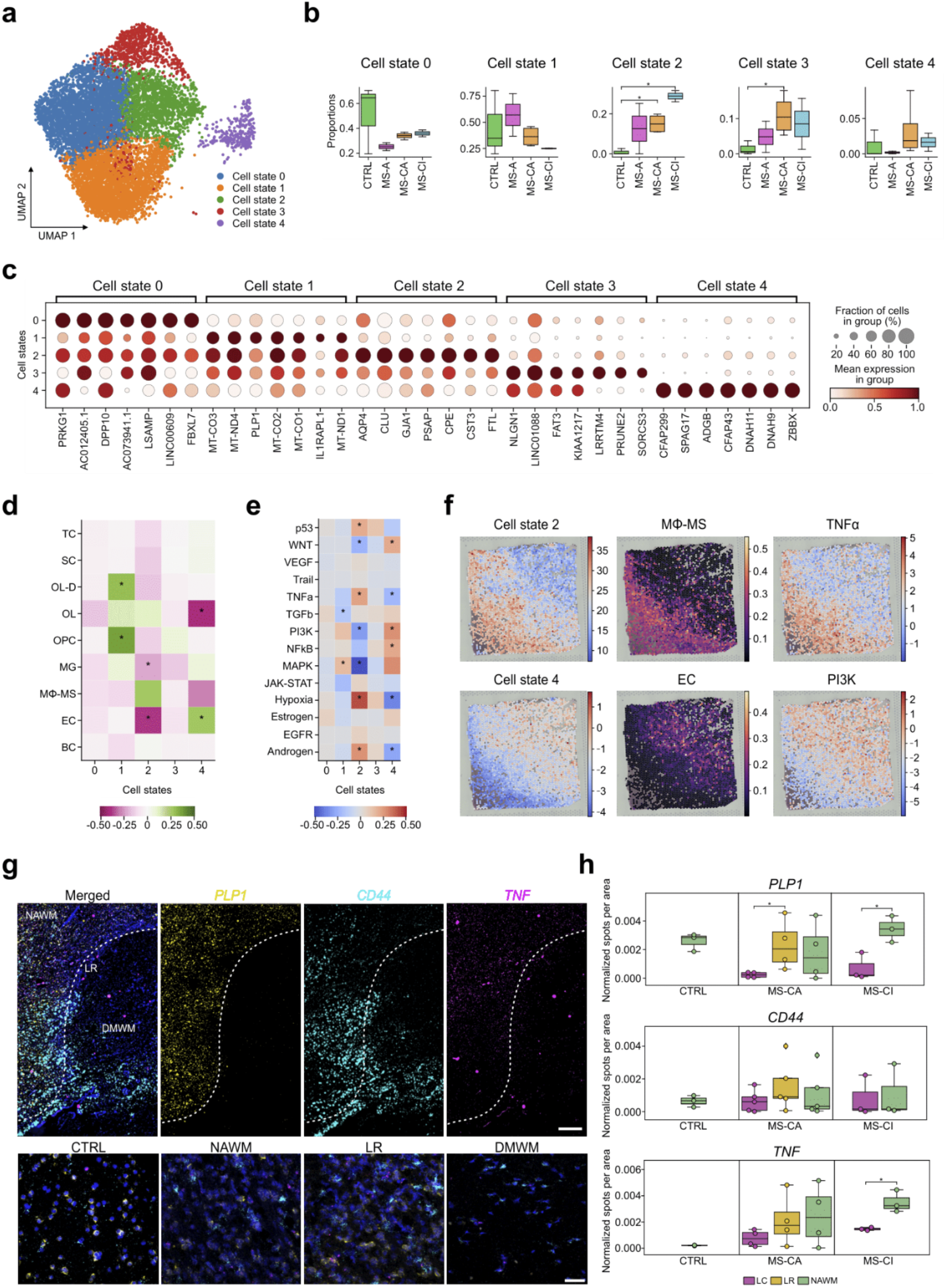
Identification and spatial characterization of AS cell states. **a**, UMAP of AS cell states based on snRNA-seq. Color indicates assigned AS subtype state. **b**, Boxplots of cell state compositions based on snRNA-seq. Color indicates MS lesion type and asterisks differences between groups (Wilcoxon rank-sum test, adjusted P < 0.15). **c**, Dotplot showing scaled mean expression of marker genes per AS cell state. **d**, Mean correlations between AS cell state scores to cell-type compositions across spots from chronic active slides. Asterisks indicate significant mean correlations (abs(mean corr) > 0.15 and mean P < 0.05). **e**, Mean correlations between AS cell state scores and pathway activities across spots from MS-CA tissue slides. Asterisks indicate significant mean correlations (abs(mean corr) > 0.15 and mean P < 0.05). **f**, Representative MS-CA tissue showing spatial colocalization of AS cell state 2 with MΦ-MS cell type mapping and TNFα activity (upper panel), and AS cell state 3 with EC cell type composition and PI3K activity. Spots with less than 11.11% of AS are removed. **g**, smFISH of MS-CA tissue for *PLP1, CD44* and *TNF*. Overview images: scale bar 500 µm; zoom-in images: scale bar 40 µm. **h**, Boxplots of smFISH quantification across different tissue areas. Color indicates different lesion and non-lesion areas, asterisks indicate differences between groups (Wilcoxon rank-sum test, adjusted P < 0.05).

Next, we profiled AS cell states by marker gene expression per cell state (Methods) (**Fig. 4c and Supplementary Table 12**). Genes associated with tissue homeostasis (*PRKG1, DPP10*)^42^ were identified in cell state 0, notably, a subtype enriched in control white matter tissues. In cell state 1, we found an enrichment of mitochondrial and myelin-related genes like *PLP1*. Of note, AS cell state 1 was associated with highly inflamed MS-A tissues characterized by ongoing myelin phagocytosis as described for reactive AS subtypes^43^. AS cell state 2 was characterized by enrichment of cell stress and reactive glial marker genes such as *AQP4, CLU* and *FTL*. Of note, increased AQP4 immunoreactivity is a key feature of reactive astrogliosis and seen in various neuropathological conditions, however, reduced in neuromyelitis optica^44,45^. Marker genes enriched in AS cell state 3 were linked to homeostatic and neuron-supporting cell functions (*NLGN1* and *LRRTM4*)^46,47^. Finally, we found that AS cell state 4 was enriched for genes encoding proteins involved in primary cilia formation (*CFAP299, SPAG17*)^48^, hence refers to the above described AS-C subtype based on snRNA-seq clustering.

To further characterize AS subtypes and their various roles in MS, we focused on MS-CA tissue samples to distinguish homeostatic from reactive subtypes in NAWM and DMWM areas, respectively. After mapping of AS states in MS-CA tissue niches (Methods), we observed that only cell states 2 and 4 showed significant correlations with other cell type abundances (P < 0.05) (**Fig. 4d**), colocalizing with MΦ-MS and EC cells respectively. Inferred pathway activity analysis (Methods) revealed that AS cell states 2 and 4, resembling glial scar and ciliated AS subtypes, correlated with both proinflammatory TNFα and tissue regeneration pathways such as PI3K and WNT, respectively (**Fig. 4e**). Next, we performed gene set enrichment analysis (Methods) and, in particular, found that AS cell state 2 was associated with lipoprotein metabolism and clearance as well as unfolded protein response; as expected, AS cell state 4 was linked to cilium assembly function (**Extended Data Fig. 3l**). Then, we mapped AS cell states to ST slides and observed that AS cell state 2 was enriched at inflamed LR areas together with MΦ-MS myeloid cells and associated with proinflammatory TNFα activity. Conversely, the ciliated AS cell state 4 mapped to DMWM areas together with EC cells resembling hypervascularized glial scar areas associated with enhanced PI3K activity (**Fig. 4f**).

To validate and map AS cell state marker genes to specific MS tissue niches, we performed smFISH for *PLP1* (to identify lesion areas), *CD44*, a canonical white matter AS marker^5^, and *TNF* encoding TNFα (Methods).

Overall, we found that *TNF* expression was upregulated in MS-CA and MS-CI relative to controls (Wilcoxon rank-sum test, adjusted P < 0.20). In detail, we observed a trend that *TNF* transcripts gradually increased at both LR and NAWM areas in MS-CA and, significantly, in MS-CI tissues (Wilcoxon rank-sum test, adjusted P < 0.05) (**Fig. 4g-h and Supplementary Tables 10-11**).

To summarize, AS subclustering and cell state-specific gene expression as well as signaling pathway analysis enabled us to identify functionally relevant disease subtypes present at distinct lesion areas as shown by spatial RNA mapping.

## Discussion

Previous tools used to assess and analyze MS pathology focused either on conventional histopathological techniques^13,14,27^ or, more recently, multiplex smFISH and snRNA-seq^5,49,1,8^ to understand cell type-specific gene expression and dysregulation. The latter tools allow for either spatial mapping of certain aspects of pathology (smFISH, histopathology) or allow cell type-specific unbiased transcriptomics (snRNA-seq). A combination of both would thus enable us to study spatially and functionally relevant signaling pathways associated with certain reactive cell types or inflamed tissue areas in the context of MS and beyond. Hence, we generated paired snRNA-seq and spatially resolved gene expression to overcome previous challenges with respect to cell type tissue profiling. Computational integration of these data types enabled us to interrogate transcriptomic profiles from cell types and their spatial location to identify patterns of cell-cell interaction and cell type cluster enrichment in specific tissue niches.

Specifically, we aimed at investigating tissue niche-resolved cell type diversity and cell-cell-communication signatures in subcortical MS tissues harboring lesions of various inflammatory activities (reflecting ‘temporal’ stages of lesion development). We generated a large-area transcriptomic map of subcortical white matter comprising control and MS donor tissues using snRNA-seq to estimate the cell type composition of tissue spots. To confirm that the computational analysis of gene expression reflects the underlying pathobiology, we compared the estimated spatial cell type maps with conventional histopathology and found a broad overlap with respect to lesion and non-lesion areas as well as myeloid cell activation and presence of iron enrichment.

In particular, we found mapping of MG/MΦ-MS subtypes (MG cell states 2 and 3) to inflamed MS-CA LR areas and mapping of AS cells including a novel subtype of ciliated AS (AS cell state 4) to demyelinated scar tissue areas (DMWM), respectively. Further, we found spatial colocalizations of certain cell types, such as SC, EC and immune cell subtypes (TC and BC) and upregulation of TGFβ signaling related to the MS-specific tissue niche 6, suggesting an increase in blood vessel density and expansion of perivascular spaces in MS. These findings would be compatible with aspects of chronic perivascular immune cell cuffing and fibrosis, indicating blood-brain impairment, and tissue inflammation^4,50,29^. Also, it is known that sustained blood-brain barrier leakage and fibrinogen deposition are pathological features of MS and experimental autoimmune encephalomyelitis^51,52^, which itself can trigger perivascular MG activation^52^. Thus, our analysis sheds light on perivascular spaces as a critical tissue niche in MS pathology. Along these lines, we demonstrate colocalization of SC and immune cells. This scenario confirms earlier data using qPCR identifying strong upregulation of genes encoding extracellular matrix proteins such as collagen chains that could be mapped to perivascular spaces in association with immune cells^29^.

Moreover, we computationally estimated signaling pathway activities associated with both tissue niches and specific cell types, such as enhanced TNFα signaling in inflamed MS-A lesion areas, characterized by MG/MΦ-MS and AS subtype enrichment, or WNT signaling linked to DMWM areas. Indeed, with respect to AS function, we observed opposing signaling pathway signatures related to their potential roles in either tissue remodeling (PI3K and WNT) or injury (TNFα). Notably, increased levels of MG-derived TNFα can trigger transformation of homeostatic AS into neurotoxic subtypes^53^, which can be facilitated by co-presence of C1q, a critical complement factor known to be upregulated on RNA and protein level in MG/MΦ-MS cells in subcortical MS lesion tissues as shown in this study and previous work^8^. Indeed, we found that both AS (cell state 2) and MG/MΦ-MS subtypes mapped to inflamed LR areas in MS-A and MS-CA tissues, suggesting functional cell-cell interactions, potentially mediated by TNFα signaling.

We found hypoxia to be another key pathway upregulated in MS-specific MG (cell states 2-4) and AS (cell state 2) subtypes. Notably, presence of hypoxia-related signaling events has been described in MS lesion pathology but, in the majority, related to neuron and OL lineage dysfunction^54,55,56^. Also, it is known that glycolysis is turned on under hypoxic conditions due to impaired oxidative phosphorylation as previously shown in proinflammatory MG and AS subtypes^57^. Further, these events have been shown to be associated with proinflammatory MG function and MG iron retention in rodent models of Alzheimer’s disease (AD), which would be in line with our findings related to proinflammatory and iron-metabolizing signatures of MS-specific MG/MΦ-MS subtypes (MG cell states 2 and 3)^58,59^.

Focusing on cell-cell communication, ligand-receptor analysis suggested that APOE-TREM2 signaling was highly specific to MG and MΦ-MS subtypes at inflamed MS-CA LR areas^8^. Indeed, enhanced APOE-TREM2 pathway activity has been implicated in other neuropathological conditions, such as AD, and associated with amyloid-beta uptake and a dysfunctional phenotype^36,60,61^. Indeed, these disease-associated MG cells might adopt senescent signatures in their ‘sustained’ attempt to clear myelin debris in MS^62^ similar to their role in the clearance of neurofibrillary tangles in tau-mediated neuropathology^62,63,64^.

Hence, these findings and predicted pathway activities illustrate how computational methods can be used to measure disease-specific pathway activity and, eventually, link these patterns to specific cell types and tissue niches in MS.

In summary, we generated and computationally analyzed a paired single-cell and spatial transcriptomics data set that enabled us to deconvolute the complex tissue microenvironment underlying lesion progression in MS by identifying both cell type-specific and spatially restricted drivers of pathology. Further, these analyses helped us better understand the molecular organization of subcortical tissues across MS lesion types. Specifically, we identified inflammatory and tissue remodeling processes that characterize subcortical lesions relative to normal white matter. As our approach is highly driven by computational prediction models in combination with histological and *in situ* validation, future work will be necessary to functionally validate the obtained findings^65^. Also, other current and future developments such as spatial ATAC-seq or ST with (sub)cellular resolution will help provide additional insight into the complex disease microenvironment in MS and beyond. However, as our results resemble important aspects and concepts from previous work, we are optimistic that our study helps break ground for novel cell type-specific therapeutic approaches to resolve compartmentalized inflammation in MS.

## Methods

### Postmortem human tissue samples

In total we examined 21 snap-frozen tissue blocks, obtained from autopsies from 16 individuals diagnosed with MS and 6 tissue donors without recognizable neuropathological changes (controls). All tissue used in this work was provided by the UK Multiple Sclerosis Tissue Bank at Imperial college in London, after ethical approval by the National Research Ethics Committee in the UK (08/MRE09/31). Further clinical and pathological information of the donors is provided in Supplementary Table 1.

### Immunohistochemistry (IHC) and histochemistry

Snap-frozen tissue blocks were dissected into 16-µm-thick slices using a Leica Microsystems CM3050S cryostat, placed on VWR superfrost plus microscope slides and stored at -80°C. Histopathological assessment was performed using immunohistochemistry for CD68 and MOG as described previously^5^ as well as CD163. The following antibodies were used: mouse anti-MOG (MAB5680, 1:1,000, Millipore Sigma), mouse anti-CD68 (MCA1815, 1:200, Bio-Rad), and mouse anti-CD163 (NCL-L-CD163, 1:1,000, Novocastra).

For CD163 immunohistochemistry and histochemistry, tissue sections were fixated on the slides by thawing and drying, followed by immersion in 4C° acetone for 10 minutes. Afterwards, the slides were allowed to dry at room temperature. Sections were stained for myelin with Luxol fast blue (LFB) by incubation with 0.1 % LFB at 56 C° overnight. After washing with 96 % ethanol and rehydration, the slides were immersed in 0.1 % aqueous lithium-carbonate solution for 5 minutes. The staining was differentiated in 70 % ethanol until the myelin sheaths obtained an intense blue color. Subsequently, tissue slides were washed with distilled water, counterstained with periodic acid-Schiff (PAS) and, following dehydration, coverslipped with a mounting medium (03989, Merck).

CD163 was stained with an Autostainer Link 48 by Dako. Endogenous peroxidase was blocked using a ready-to-use peroxidase-blocking solution (S202386-2, Dako). The primary antibody was diluted in antibody diluent (S080983-2, Dako) and applied for 60 minutes at room temperature (RT). After a washing step, the slides were exposed to a mouse-specific biotinylated secondary antibody (GV82111-2, Dako) for 15 minutes, followed by a streptavidin-linked horseradish peroxidase (SM802, Dako) for 20 minutes. The staining was developed using a 1:50 dilution of 3,3’-diaminobenzidine (DAB) chromogen in DAB+ substrate buffer (GV825, Dako) for 10 minutes. Sections were counterstained using hemalaun and coverslipped.

To localize ferrous and ferric iron, DAB-enhanced Turnbull Blue (TBB) was performed as previously described in another study^62^. In short, the slides were dried and exposed to 10% ammonium sulfide solution (105442, Merck) in double-distilled water for 90 minutes. Consecutively, slides were immersed in 10% potassium ferricyanide and 0.5% hydrogen chloride in an aqueous solution for 15 minutes. This step was followed by blocking the endogenous peroxidase with 0.01M sodium azide and 0.3% hydrogen peroxide in methanol for 60 minutes. The slides were washed with 0.1 M Sorensen’s phosphate buffer and the staining was developed with a 1:50 solution of DAB chromogen (K3468, Dako) and 0.005% hydrogen peroxide in Sorensen’s phosphate buffer for 20 minutes. Slides were counterstained with hemalaun and coverslipped.

### Brightfield microscopy and image processing

Brightfield images of LFB, CD163 and TBB were acquired using a Leica DMi8 microscope and a Hamamatsu NanoZoomer 2.0HT. Images were taken at a 40x magnification and exported as NPD files. Image processing of histological data was performed using GIMP-2.10 software.

### Histopathological assessment and lesion type characterization

In our analyses, only white matter lesions were selected. Lesions were identified as areas with a marked loss of myelin by anti-MOG and LFB-PAS staining, and further classified into MS-A, MS-CA and MS-CI typed lesions using CD68 and CD163 immunohistochemistry for MG and MΦ-MS subtypes according to established protocols and criteria^15^. Of note, no differentiation was made between early and late active-demyelinating MS-A lesions. Fully remyelinated lesions, so-called shadow plaques, were not included.

The lesion area was subdivided into distinct zones. The lesion center or core covers the middle of the demyelinated lesion towards the rim, which is the distinct border between demyelinated lesion center and the surrounding myelinated white matter, called peri-plaque white matter (PPWM). The PPWM covers the surrounding white matter and transitions at a distance of one centimeter from the lesion border into the normal-appearing white matter (NAWM).

The lesion types were assigned by a trained neuropathologist. Acute lesions were identified by a hypercellular lesion center and a high density of CD68- and CD163-positive macrophages and microglia, also containing LFB- and/or MOG-positive myelin degradation products, throughout the lesion, and an indistinct LR. Chronic active lesions showed a hypocellular, demyelinated lesion center but a distinct rim formation by CD68- and CD163-positive cells, containing LFB-positive myelin degradation products. Some of these lesions showed accumulation of iron-laden microglia and macrophages at the rim. Chronic inactive lesions were characterized by a fully demyelinated, hypocellular lesion center, and a low frequency of CD68- and CD163-positive macrophages or microglia within the lesion, and a distinct LR without accumulation of CD68- or CD163-positive cells.

### Sample selection for transcriptomics

The RNA integration number (RIN) was used as a sample selection criteria, and only samples with a value of ≥ 6 were included for both transcriptomic analysis. We cut 70µm thick sections of tissue on a CM3050S (Leica) to obtain a final weight of 15mg of tissue, from which the RNA was isolated. This was done using TRIzol (15596026, Thermo Fisher), chloroform (1731042, Sigma Aldrich) and RNAeasy mini Kit (74104, QIAGEN) following manufacturer’s recommendations. RNA integrity was measured on an Agilent 2100 Bioanalyzer using the High Sensitivity ScreenTape (5067-5579, Agilent), buffer (5067-5580, Agilent) and ladder (5067-5581, Agilent) according to manufacturer’s instructions.

### Nuclei isolation and libraries preparation

Nuclei from selected samples were isolated using a sucrose-gradient ultracentrifugation according to established workflows^5^. Following isolation, nuclei were diluted to a final concentration of 1000 nuclei per µl and loaded to the 10x Genomics Chromium controller aiming for a recovery rate of 8000 nuclei per sample. We prepared the libraries following the 10X Genomics protocol using the 3’ single cell v3.1 kit (PN 1000121) with single indexing. Samples were sequenced on a NovaSeq 6000 aiming for a sequencing depth of 30000 reads per nuclei. Expression count matrices for each sample were generated using Cell Ranger Count v. 6.0.2 by performing alignment to the sequencing data against the GRCh38-2020-A reference transcriptome.

### snRNA-seq data quality control

The data processing and downstream analyses for the snRNA-seq datasets was done using the scanpy toolkit (v1.8.2)^66^ in Python (v.3.9.12). Each sample was filtered separately to control for batch differences. Single nuclei were filtered by genes (200 < number of genes), mitochondrial genes (percentage of mitochondrial genes < 5%) and gene counts (number counts < number counts 99th percentile). Genes were kept if they were expressed across different nuclei (number of expressed nuclei > 3). Dissociation scores (score_genes) were computed with score_genes using the top 200 genes ranked by p-value from a list of genes previously associated with tissue dissociation and cell death^67^ and doublet scores were computed using scrublet^68^. Afterward, nuclei were filtered by dissociation scores (dissociation score < dissociation score 99th percentile), and doublet scores (doublet score < 0.2). Finally, each nuclei raw expression was normalized by a total sum of 10,000 and log-transformed (log1p).

### snRNA-seq data integration and cell annotation

A single AnnData object was generated by concatenating (join=“outer”) all the preprocessed nuclei coming from different samples. Feature selection was performed by computing high variable genes per sample and then selecting the top 3,000 genes that were flagged as variable in the maximum number of samples. Genes were then scaled across nuclei (max_value=10) and PCA was calculated on the selected features. Harmony-py^69^ was used to integrate the obtained PCs, eliminating batch effects between samples. Nearest neighbors were generated for nuclei by estimating similarities in the PC space (n_pcs = 50). The obtained connectivities were used to generate a UMAP manifold. Nuclei were clustered using the Leiden graph-clustering method^70^ (resolution = 0.5) and annotated manually using brain gene markers^58^. Clusters belonging to gray matter cells or that showed no clear cell-specific markers were removed from downstream analyses. After removing cell clusters, the steps of highly variable gene selection, integration of nuclei and annotation of clusters were repeated until no more clusters had to be removed.

### Comparison with an independent atlas

To validate our cell type annotations, we compared our generated atlas with another reference human snRNA-seq atlas^8^ at the molecular level. The counts matrix and annotation meta-data were downloaded from https://www.ncbi.nlm.nih.gov/geo/query/acc.cgi?acc=GSE180759. Pseudo-bulk transcriptomic profiles were generated for both atlases at the cell type level using decoupler-py (v1.2.0)^71^ with the following hyperparameters: groups_col=None, min_prop=0.2; min_smpls=0. To make them comparable, we filtered the profiles by the intersection of genes between the two atlases and log normalized them with scanpy^66^. Finally, the Pearson correlation between the different profiles was performed.

### ST workflow

10x Genomics Visium Spatial Gene Expression platform was used for the spatial transcriptomics experiments. The tissue (RIN≥7) was cut into 10 µm sections using Leica CM3050 S cryostat and placed into a Spatial Gene Expression Slides (PN-1000185, 6,5 × 6,5 mm ROI) that was precooled inside the cryostat at -22°C. The slides were stored in a container at -80°C until further processing. The sections were then fixed and stained using protocol CG000160 Rev B. The sections were then imaged, to do a general morphological analysis and for future spatial alignment of the data, using the 10x brightfield objective from the Leica DMi8 and processed by the Leica Application Suite X (LAS X).

Then they were enzymatically permeabilized for 18 min. This time was assessed using the 10x Visium Tissue Optimization kit (PN-1000191) and following the protocol CG000238 Rev D.

The generation of the libraries was performed according to published protocols (10x Genomics): CG000239 Rev D, using the Gene Expression Reagent kit (PN-1000186), the Library Construction Kit (PN-1000190) and the Dual Index Plate TT Set A (PN-1000215). In order to assess the correct amplification of obtained cDNA, QuantStudio 3 from ThermoFisher was used. For full length of cDNA and indexed libraries analysis, the TapeStation 4200 analyzer (Agilent) was used. The libraries were loaded at 300 pM and sequenced on a NovaSeq 6000 system (Illumina) with a sequencing depth of 250 million reads per sample.

The demultiplexing of the data was done using SpaceRanger software (10x – version 1.2.2), creating FASTQ files. These files are then used by SpaceRanger count to perform alignment with the human reference genome GRCh38-2020, tissue detection, fiducial detection and barcode/UMI counting.

### ST data quality control

First, spatial coordinates belonging to cortical gray matter areas were manually removed using the Loupe software (10x Genomics). Then, the following data processing and downstream analyses for the ST datasets were performed using the scanpy toolkit^66^. For each slide, spots were filtered by genes (200 < number of genes) and genes were kept if they were expressed across different spots (number of expressed spots > 3).

### ST cell type deconvolution

The cell2location (v0.1)^72^ package was used to calculate cell type abundances for each spot. From the annotated snRNA-seq atlas, reference expression signatures of major cell types were inferred leveraging regularized negative binomial regressions. Then each slide was deconvoluted using hierarchical bayesian models with the following hyperparameters: N_cells_per_location=5 and detection_alpha=20. Afterwards, cell type proportions were calculated per spot by dividing the abundance of a given cell type by the total sum of abundances of a given spot. To assess the quality of the deconvolution, we calculated the Pearson correlation between the mean estimated cell type proportions of each slide with the observed cell type proportions in its corresponding snRNA-seq dataset.

### Generation of cell type compositional niches in ST

Cell type compositional niches were generated from the estimated cell type proportions in ST slides. First, cell type proportions from ST slides were forced to sum to one, and then they were transformed using the isometric log ratio transformation^73^ with the composition-stats python package (v2.0.0). Then, these features were integrated across slides using harmony-py^69^ correcting for technical effects. Afterwards, nearest neighbors (n_pcs = 50) were computed to generate a UMAP manifold. Finally, spots were clustered using the Leiden graph-clustering method^70^ (resolution = 0.4). Clusters that were overrepresented by only one sample were removed.

### Characterization of ST niches

We determined representative cell types for each ST niche by first computing the mean proportion of each cell type per slide, obtaining a distribution of correlations, and then computing the mean for each niche. Then, to compare cell type correlations between niches we used the Wilcoxon rank-sum test (adjusted P < 0.05).

To validate these representative cell types across niches, we computed for each slide the pairwise Pearson correlation between cell types. To aggregate the results, we computed the mean correlation and p-value across slides. Interactions with abs(correlation) > 0.15 and P < 0.10 were considered as significant.

We further characterized niches by inferring signaling pathway activities for each spot with the resource PROGENy^74^ and the method Univariate Linear Model (ULM) from decoupler-py (v1.2.0)^71^. We selected the top 300 most significant genes per pathway from PROGENy. Then we tested which pathways are representative for each niche by computing the mean activity of each pathway per niche and slide. Then, we calculated the mean of each niche across slides and compared them using the Wilcoxon rank-sum test (FDR < 0.05).

### Comparison of tissue architecture across lesion types

To identify differences between lesion types, we first compared them at the transcriptomic level. Global pseudo-bulk transcriptomic profiles for snRNA-seq samples and ST slides were generated using decoupler-py (v1.2.0)^71^. Next, counts were aggregated for each sample/slide for genes that were expressed in at least 20% of cells/spots and a minimum of three samples/slides with the get_pseudobulk function (groups_col=None). Then each raw expression profile was normalized by a total sum of 10,000 and log-transformed (log1p). Afterwards, profiles were scaled by genes and PCA (PCs=50) was performed.

Mean cell type proportions across snRNA-seq samples and ST slides were compared with Kruskal-Wallis tests (adjusted P < 0.075). For the significant cell types, pairwise comparisons were performed with the Wilcoxon rank-sum test (FDR < 0.1). Additionally, we performed Kruskal-Wallis tests over the compositions of niches (adjusted P < 0.05) and the corresponding pairwise comparisons using the Wilcoxon rank-sum test (FDR < 0.1).

To identify cell types that colocalize differentially across lesion types, we computed the correlation between cell types per sample. Then, we compared the obtained distributions of correlations with the Kruskal-Wallis tests (adjusted P < 0.1) and for the significant interactions we performed pairwise comparisons between lesion types using the Wilcoxon rank-sum test (adjusted P < 0.1).

### Molecular differences between chronic active and control in snRNA-seq

To identify molecular differences between control and MS-CA tissue samples in snRNA-seq, we first generated log-normalized pseudo-bulk transcriptomic profiles for each sample and cell type using decoupler-py (v1.2.0)^71^ and computed differential genes between lesions using the t-test with BH correction for the obtained p-values. Afterwards, mitochondrial genes, genes with no significant change (abs(t-statistic) < 0.5 and P > 0.05) and genes that were significant across multiple cell types were removed.

From the positively regulated differential expressed genes in chronic active, we inferred transcription factor and pathway activities from their t-statistics. For transcription factors, we leveraged DoRothEA^75^, a gene regulatory network generated from prior knowledge, and the method multivariate linear model from decoupler-py (v1.2.0)^71^. For pathways we used the resource PROGENy^74^ and the method Univariate Linear Model (ULM) from decoupler-py (v1.2.0)^71^.

### Identification of cell states

To identify cell states for MG and AS cells, we subsetted the corresponding annotated cells from the complete snRNA-seq atlas. For AS, we selected both AS and AS-C, the following described steps were performed separately for MG and AS. We removed genes that were not expressed in at least three nuclei and log normalized the read counts. We applied feature selection by computing high variable genes per sample and then selecting the top 3,000 genes that were flagged as variable in the maximum number of samples. Afterwards, genes were scaled across nuclei (max_value=10), PCs were calculated on the selected features (PCs = 50) and they were integrated using harmony-py^69^. Finally, nearest neighbors (n_pcs = 50), UMAP manifold and clustering of nuclei with the Leiden graph-clustering method^70^ (resolution = 0.5) were performed. Clusters that were overrepresented by only one sample were removed.

### Identification of marker genes for cell states in snRNA-seq

Differential expression analysis between cell states in snRNA-seq was performed to identify marker genes using the rank_genes_groups (method=t-test_overestim_var) function from scanpy with the log normalized counts. Genes with abs(logFC) > 1 and adjusted P < 0.05 were considered marker genes for a given cell state.

### Cell-state spatial mapping

To map cell states to spatial locations, we used leveraged deconvolution results and a set of marker genes of each cell state. For a given major cell type, we selected the spots which hold its proportion to at least 11.11% and inferred its cell state activities per spot using previously obtained marker genes as a resource combined with the method ULM from decoupler-py (v1.2.0)^71^.

### Inference of activities for biological terms in cell-states

For the inference of biological processes, we used gene sets extracted for the resource MSigDB^76^ with the ULM method from decoupler-py (v1.2.0)^71^. Enrichment activities were inferred per nucleus, and then the mean scaled activity per cell state was calculated.

### Inference of cell-cell communication events

We inferred ligand-receptor interactions between cell states of microglia and other relevant cell types from snRNA-seq in chronic active samples using the LIANA(v0.1.6) framework^77^. We inferred interactions using the default cell-cell communication methods provided in LIANA with OmniPath^78^ as prior knowledge resource and selected the ones found to be significant after performing the consensus ranking between methods (P < 0.05). Next, we checked whether the found interactions exist in our ST data by computing the correlation between the gene expression of a given ligand with a given receptor when both cell types are present in the same spot with a proportion > 0.11.

### Fluorescence multiplex in situ RNA hybridization

For single-molecule fluorescence *in situ* hybridization (smFISH) validation, we used frozen human cryosections of 16µm thickness. smFISH was performed on a representative selection of tissue samples using ACD RNAscope 2.5 HD Red and Multiplex Fluorescent V2 assays (ACD Biotecne), following the protocol of a previous publication^1^. The following human RNAscope assay probes were used: *FTL* (C1), *PLP1* (C1), *APOE* (C2), *CD44* (C2), *TREM2* (C3) and *TNF* (C3).

### Image acquisition and quantification of multiplexed fluorescent images

Overview and quantification images were taken using a Leica DMi 8 microscope with a Leica DFC7000 GT camera. Focus points were set at 10x magnification for overview purposes and at 20x magnification at the area of interest for quantification. Pictures were imaged and exported as LIF files. A Leica TCS SP8 microscope was used for taking confocal images. All fluorescent pictures are z-stack images consisting of 10 to 20 layers with a 0.5–0.7 μm step size. Heights for z-stack were identified manually by imaging DAPI on the area of interest. Each z plane was imaged across 4 channels.

To detect spatial alterations in expression profiles throughout the lesion area, we defined four regions of interest (ROIs) that were examined in every lesion. The following ROIs were considered: normal white matter from controls (NWM), normal appearing white matter (NAWM) of MS patients in at least 1000µm distance from the lesion rim (LR) that was defined as the region directly bordering DMWM areas with a thickness of 400µm. Lesion center (LC) was defined as an area with a distance of 500-1000µm to the LR.

Single-molecule FISH (smFISH) data was analyzed using RS-FISH^79^. Multi-channel images were split into single-channel images using bfconvert from the bftools suite^80^. RS-FISH was run on each channel using the command-line implementation with the following parameters (channel 1 (*FTL, PLP1*): sigma = 1.43, threshold = 0.006, channel 2 (*APOE, CD44*): sigma = 1.4, threshold = 0.012, channel 3 (*TREM2, TNF*): sigma = 1.46, threshold = 0.006) and ransac=1. The resulting spot count tables from RS-FISH were assigned to manually annotated mask regions of the image. Regions of interest (ROI) were drawn on images in Fiji and converted to multi-class labeled masks using the Mask(s) from ROI(s) plugin (https://imagej.net/plugins/masks-from-rois). Spot counts for each mask were subsequently normalized to the mask area. Pairwise comparisons of the normalized spots per area were performed using the Wilcoxon rank-sum test (adjusted P < 0.2).

## Acknowledgements

We thank Djordje Gveric (Imperial College London) for help with selection of human brain samples from the UK MS Tissue Bank, funded by the MS Society of Great Britain and Northern Ireland and Angelika Duda (Medical Faculty Mannheim, Heidelberg University) for technical assistance and help with snRNA-seq preparations. We gratefully acknowledge the data storage service SDS@hd and high-performance computing service bwHPC supported by the Ministry of Science, Research and the Arts Baden-Württemberg (MWK) and the German Research Foundation (DFG) through grant INST 35/1314-1 FUGG. This work was supported by intramural funding provided by the Medical Faculty Mannheim of Heidelberg University (to L.S.), a German Cancer Aid scholarship (to T.T.), research grants from the Hertie Foundation (medMS MyLab, P1180016 to L.S.), the European Research Council (‘DecOmPress’ ERC StG, to L.S.), the National Multiple Sclerosis Society (RFA-2203-39300, to L.S., PA-2002-36405, to S.H. and L.S.), the German Research Foundation (SCHI 1330/2-1, 4-1 and 6-1 to L.S., GRK2727 student fellowships to C.L.M. and P.S., WU 1098/1-1 Walter-Benjamin position to F.W.), the German Federal Ministry of Education and Research (BMBF 01ZZ2004, to F.W., M.I. and D.S.) the Silicon Valley Community Foundation (2017-171531(5022) to M.H.) and the National Human Genome Research Institute and NIH (5U41HG002371-23 to M.H.). This work was supported by the DFG Research Infrastructure NGS_CC (project 407495230) as part of the Next Generation Sequencing Competence Network (project 423957469). NGS analyses were carried out at the Competence Center for Genomic Analysis (Kiel, Germany).

## Author contributions

L.S. and J.S.R. conceived the study. C.L.M. and A.H. selected and assessed tissue samples from control and MS donors (advised by L.S.). A.H. and C.J.R. acquired images for histopathological assessment. C.L.M., A.H. and C.J.R. classified lesions (advised by L.S., P.E. and S.H.). C.L.M. and T.H.T. performed nuclei isolation.

C.L.M. carried out the snRNA-seq libraries and ST experiments. C.L.M. and A.H. performed smFISH. A.H., T.H.T. and T.T. acquired images for smFISH quantification. P.B.M. and R.O.R. designed the computational analysis (supervised by J.S.R.). P.B.M. analyzed and integrated snRNA-seq and ST datasets (advised by C.L.M., R.O.R., J.S.R. and L.S.). F.W. and M.I.A. performed smFISH image analysis (supervised by D.S.). M.H. generated the web browser for visualization of snRNA-seq and ST data sets. C.L.M., P.B.M., J.S.R. and L.S. wrote the manuscript. C.L.M., P.B.M., P.S., and L.S. organized figures. All authors have read, edited and approved submission of the manuscript.

## Data and code availability

Data will be made available to the public after peer review. The custom scripts used in this work are available on Github: https://github.com/saezlab/VisiumMS.

---

**Extended Data Fig. 1.**
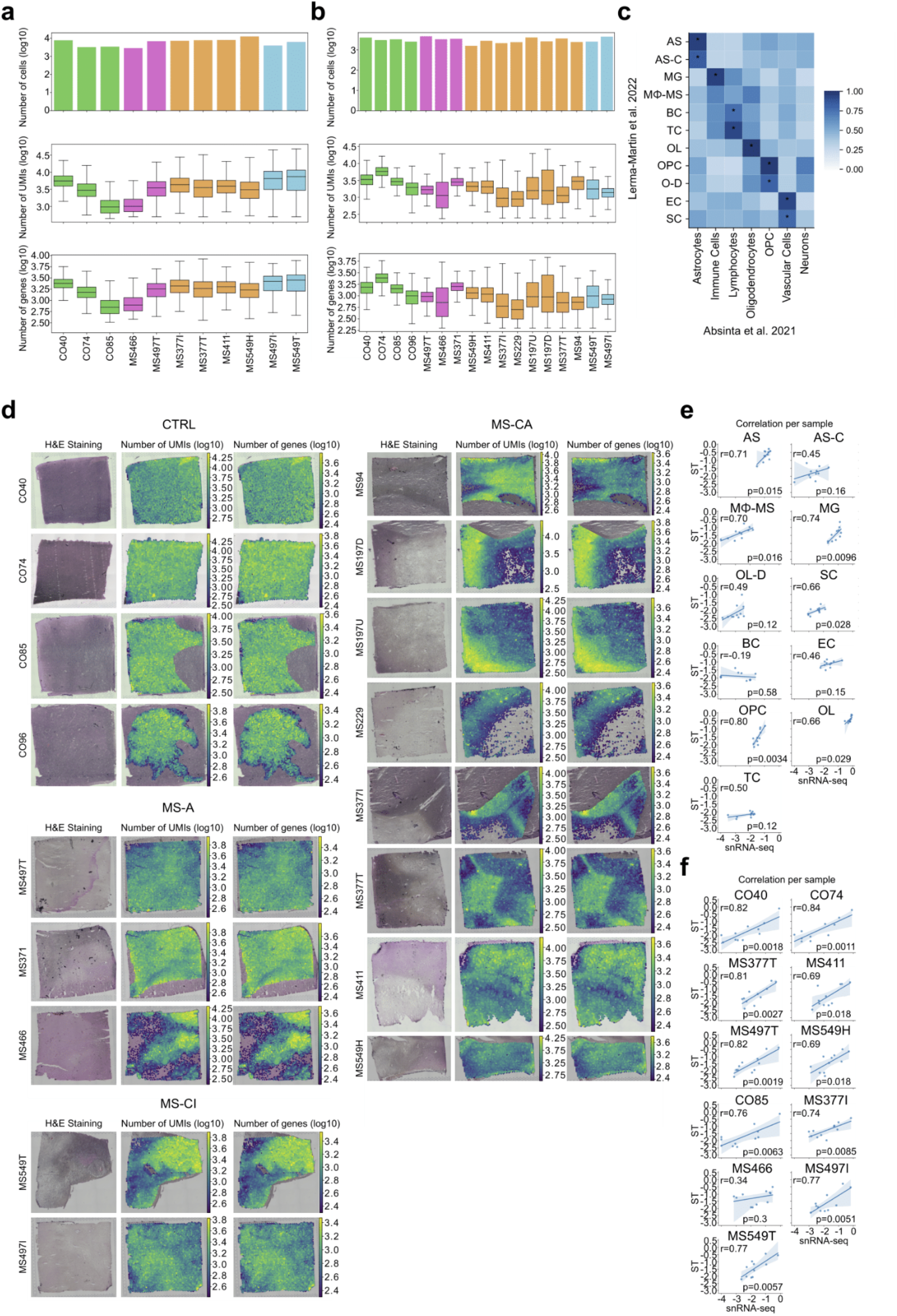
Subcortical snRNA-seq and ST quality metrics. **a**, Quality control metrics for snRNA-seq samples. **b**, Quality control metrics for ST samples. **c**, Comparison of subcortical snRNA-seq atlas with previously published subcortical MS atlas according to Absinta et al., 2019. Color intensity indicates Pearson correlation between cross-study snRNA-seq cell type profiles, and stars indicate significant interactions (P < 0.05). **d**, HE tissue histology and quality control metrics per ST spot across tissue slides. **e**, Pearson correlation between snRNA-seq cell type proportions and obtained proportions from cell type deconvolution at cell type level in log scale. **f**, and at the sample level in log scale.

**Extended Data Fig. 2.**
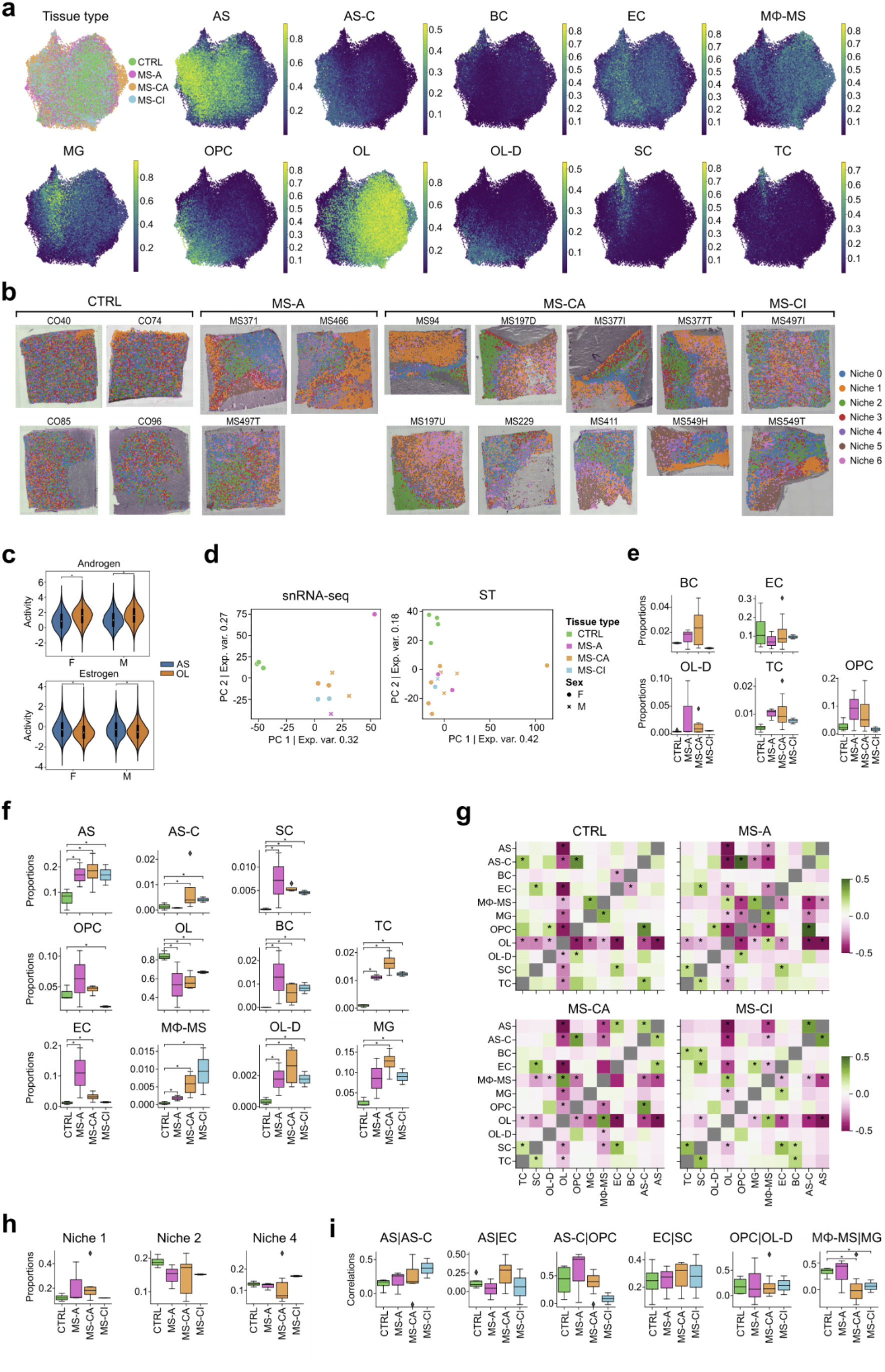
Subcortical cell type and tissue niche mapping metrics. **a**, UMAP of spatial cell type mapping based on ST spots and cell type tissue composition. Color indicates cell type composition values and MS lesion types. **b**, Visualization of obtained tissue niches across samples. Color indicates niche category. **c**, Comparison of pathway activities for Androgen and Estrogen across niches per gender, F females, M males. Color indicates AS-enriched niches (1, 5 and 6), and OL-enriched niches (0, 2, 3 and 4), asterisks indicate significance (Wilcoxon rank-sum test, P < 0.05). **d**, PCA projection of first two components of snRNA-seq and ST transcriptomic profiles. Color indicates lesion type and shape gender. **e**, Boxplots showing deconvoluted ST cell type composition per tissue type. Color indicates lesion type. **f**, Boxplots showing snRNA-seq cell type composition per tissue types. Color indicates lesion type and asterisks significant differences between groups (Wilcoxon rank-sum test, adjusted P < 0.1). **g**, Mean Pearson correlations between cell type composition across spots per lesion type. Asterisks indicate significant mean correlations (abs(mean corr) > 0.1 and mean P < 0.1). **h**, Boxplots of niche compositions across lesion types. Color indicates lesion types. **i**, Boxplots of colocalization correlations between cell types. Color indicates lesion types.

**Extended Data Fig. 3.**
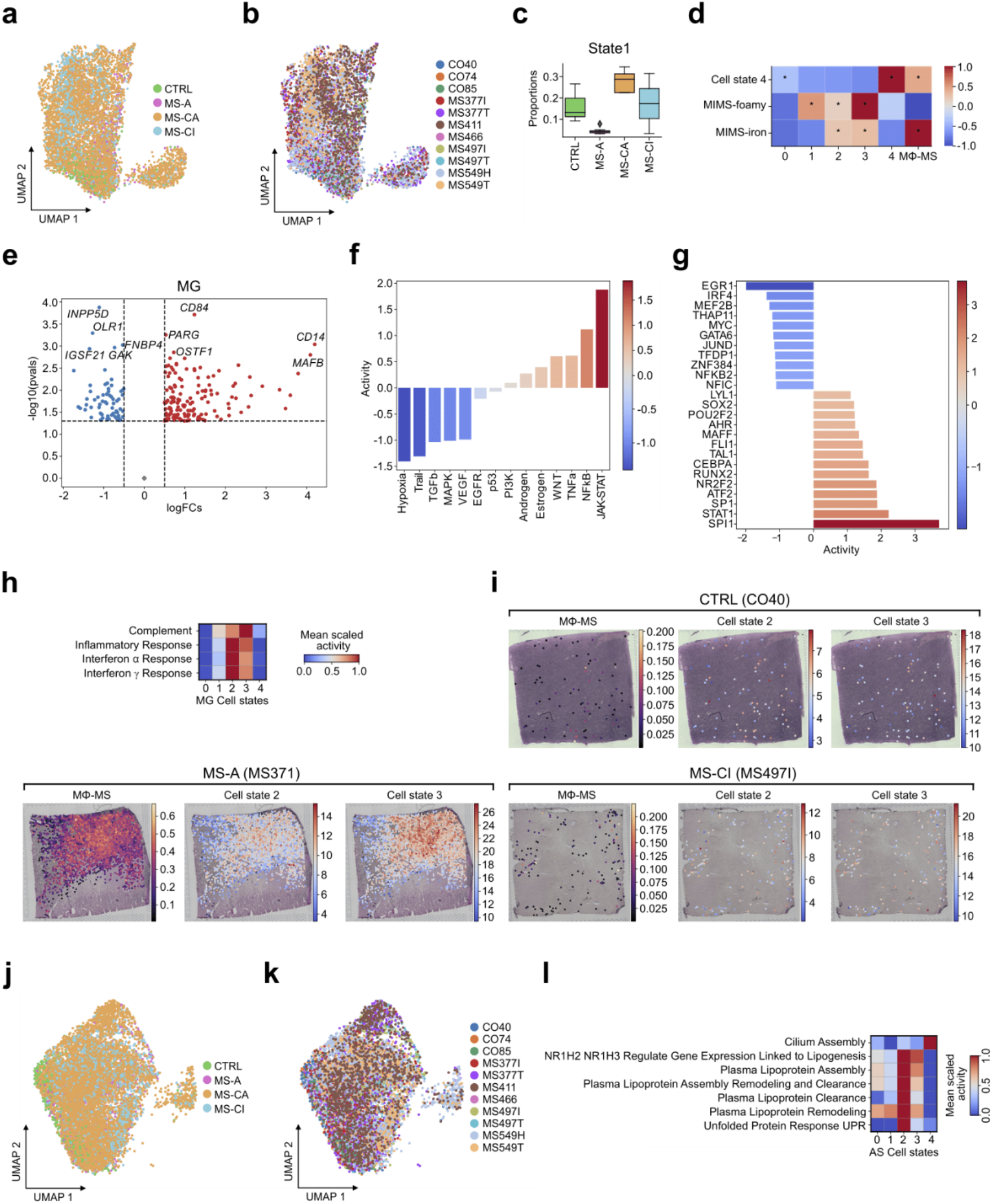
Identification and spatial characterization of MG and AS cell states. **a**, UMAP of MG cell states, color corresponds to the lesion type of the samples. **b**, UMAP of MG cell states, color corresponds to the tissue sample IDs. **c** MG cell state 1 composition per MS lesion type. **d**, Myeloid cell subtype gene expression correlation analysis between *Absinta et al* and current study. **e**, Volcano plot of differentially expressed genes in all MG cells between MS-CA and control tissue samples (abs(logFC) > 0.5 and P < 0.05). Red color indicates positively regulated genes and blue color negatively regulated ones. **f**, MG pathway activity (most upregulated [red] vs. most downregulated [blue]) in MS-CA relative to control tissue samples. **g**, MG transcription factor activity (most active [red] vs. most inactive [blue]) in MS-CA relative to control tissue samples. **h**, Activities of biological terms between MG cell states. **i**, Representative ST samples demonstrating colocalization of MΦ-MS cell type with MG states 2 and 3 across MS lesion types. Spots with less than 11.11 % of MG were removed. **j**, UMAP of AS cell states, color corresponds to the lesion type of the samples. **k**, UMAP of AS cell states, color corresponds to tissue sample IDs. **l**, Activities of biological terms between AS cell states.

